# Widespread shorter cortical adaptation in dyslexia

**DOI:** 10.1101/219923

**Authors:** Sagi Jaffe-Dax, Eva Kimel, Merav Ahissar

## Abstract

Studies of dyslexics’ performance on perceptual tasks suggest that their implicit inference of sound statistics is impaired. In a previous paper (Jaffe-Dax, Frenkel, & Ahissar, 2017), using 2-tone frequency discrimination, we found that the effect of previous trial frequencies on dyslexics’ judgments decayed faster than the effect on controls’ judgments, and that the adaptation of their ERP responses to tones recovered faster. Here, we show the cortical distribution of this abnormal dynamics of adaptation using fast acquisition fMRI. We find that dyslexics’ faster decay of adaptation is widespread, though the most significant effects are found in the left superior temporal lobe, including the auditory cortex. This broad distribution suggests that dyslexics’ faster decay of implicit memory is a general characteristic of their cortical dynamics, which also encompasses the sensory cortices.

## Introduction

Dyslexia, a specific and significant impairment in the development of reading skills that is not accounted for by mental age, visual acuity problems, or inadequate schooling (WHO, 2010), affects ˜5% of the world’s population (Lindgren, De Renzi, & Richman, 1985). Though dyslexics are diagnosed for their reading impairments, they also often have difficulties on simple non-linguistic perceptual tasks (Ahissar, Protopapas, Reid, & Merzenich, 2000; Giraud & Ramus, 2013; McAnally & Stein, 1996; Sperling, Lu, Manis, & Seidenberg, 2005). These can be largely explained as resulting from inefficient use of stimulus statistics in the experiment (the “Anchoring Deficit hypothesis”; Ahissar, Lubin, Putter-Katz, & Banai, 2006; Oganian & Ahissar, 2012, Jaffe-Dax et al., 2015). In these tasks, participants are not aware of the effect of previous stimuli, but their perception tends to contract to their estimated mean of these stimuli (contraction bias; Raviv, Ahissar, & Loewenstein, 2012; Raviv, Lieder, Loewenstein, & Ahissar, 2014).

The neural mechanism that may underlie the implicit learning of experimental statistics is adaptation; i.e., an automatic, implicit, and stimulus-specific decrease of the response to repeated stimuli. Importantly, the rate of decay of the behavioral effect of previous trials in serial discrimination is similar to the rate of decay of neural adaptation, as measured by MEG (Lu, Williamson, & Kaufman, 1992). Inspired by this finding, we recently compared both behavioral dynamics and rate of adaptation (ERP responses) of good readers (i.e., the control group) and dyslexics (Jaffe-Dax, Frenkel, & Ahissar, 2017). The participants performed serial discrimination in four blocks of trials with different Trial Onset Asynchronies (TOAs). Both the magnitude of perceptual contraction to the mean frequency of previous trials and the magnitude of neural adaptation (P2 and N1 components that are automatically produced by the auditory cortex, Mayhew, Dirckx, Niazy, Iannetti, & Wise, 2010) decayed faster in dyslexics (ERP; Jaffe-Dax et al., 2017).

Since ERP responses cannot be used to localize the cortical source of this group difference, we then recruited the participants from the ERP study (Jaffe-Dax et al., 2017) to take part in an fMRI study with a similar protocol, which allowed us to characterize which brain areas show shorter adaptation in dyslexics. Using the ERP based protocol in the scanner, we measured the BOLD response (βs) to tones for each TOA, and calculated the time constant of adaptation (fitting an exponential decay function) in the responding voxels and in the (pre-defined) auditory cortex. All cortical regions that responded to tone discrimination showed a tendency to decay faster in dyslexics. Significant differences were found in the primary auditory cortex, broader regions of the left superior temporal lobe, and in the right insular cortex.

## Results

We recruited 20 dyslexics and 19 good readers from our previous study (Jaffe-Dax et al., 2017) and asked them to perform 2-tone frequency discrimination in separate blocks with four trial-onset intervals (TOAs) of 3, 6, 9, and 15 seconds, respectively. Before entering the scanner, all participants performed a short 4-block training session, in which the two groups exhibited similar accuracy (72.4 ± 6% vs. 73 ± 4.6%, *z* = 0.5, *p* = 0.57). In-scan, good readers (controls) performed better (82.5 ± 1.6% vs. 76.3 ± 2.2%, *z* = 2.6, *p* < 0.01. Mean ± SEM. Mann-Whitney U-tests), suggesting that they gained more from the short pre-scan practice (in line with the faster learning reported in Jaffe-Dax et al., 2017).

To evaluate the dynamics of cortical adaptation in each group, we used the following procedure. First, we determined which Talairach voxels responded to the task (standard GLM, *p* < 0.001, FDR corrected) when all participants were considered. For each of these voxels, we calculated the dynamics of adaptation, among controls and among dyslexics, as follows. We estimated β over the mean BOLD response of each group in each of the four TOA conditions. Using these βs, we fitted an exponential decay model (Jaffe-Dax et al., 2017): β *TOA* = *a* + *b* exp ϕ*TOA* τ to each voxel. In this model τ denotes the time scale of adaptation, *a* is the asymptote level of BOLD and ^controls^*b* is the amplitude of adaptation. Figures 1A-B plot the distribution of the fitted τs for and dyslexics, respectively. It illustrates the broadly distributed trend of faster decay in the dyslexic group.

To locate regions in which the fitted τ differed significantly between the groups, we conducted a whole brain analysis, in which we fitted τ to each voxel, and for each participant separately. To reduce the impact of outliers resulting from the noisy estimation of τ (due to this single subject & single voxel analysis) we assessed group difference with a non-parametric test (Mann-Whitney U test), in which extreme values are not over-weighted. We corrected for multiple comparison bias by requiring a cluster of contingent voxels with a significant group difference (*p* < 0.05, cluster corrected to 44 spatially contingent voxels, based on Monte-Carlo cluster level correction). Significant regions were found in the left superior temporal cortex (TAL: −54, −18, 10) and in the right insular cortex (TAL: 39, −2, −8), outlined in purple in Figures 1A-B. The superior temporal cortex is known to be involved in a broad range of auditory tasks, including simple tone discrimination (Daikhin & Ahissar, 2015), language (Fedorenko, Hsieh, Nieto-Castañón, Whitfield-Gabrieli, & Kanwisher, 2010), music (Fedorenko, Behr, & Kanwisher, 2011), and even social tasks (e.g. Deen, Koldewyn, Kanwisher, & Saxe, 2015). Thus, this group difference for this area was expected given the behavioral results. The right insular cortex is multi-modal (Bushara et al., 2003), and is also involved in introspection (Craig, Chen, Bandy, & Reiman, 2000). Comparing Figures 1A and 1B suggests that other regions might have a larger mean group differences (e.g., frontal cortices), but due to large inter-subject variability in these regions, the group differences were not significant. This large variability might account for the spurious dots of large τ values scattered throughout the cortical map (Figure 1 A-B).

**Figure 1.**
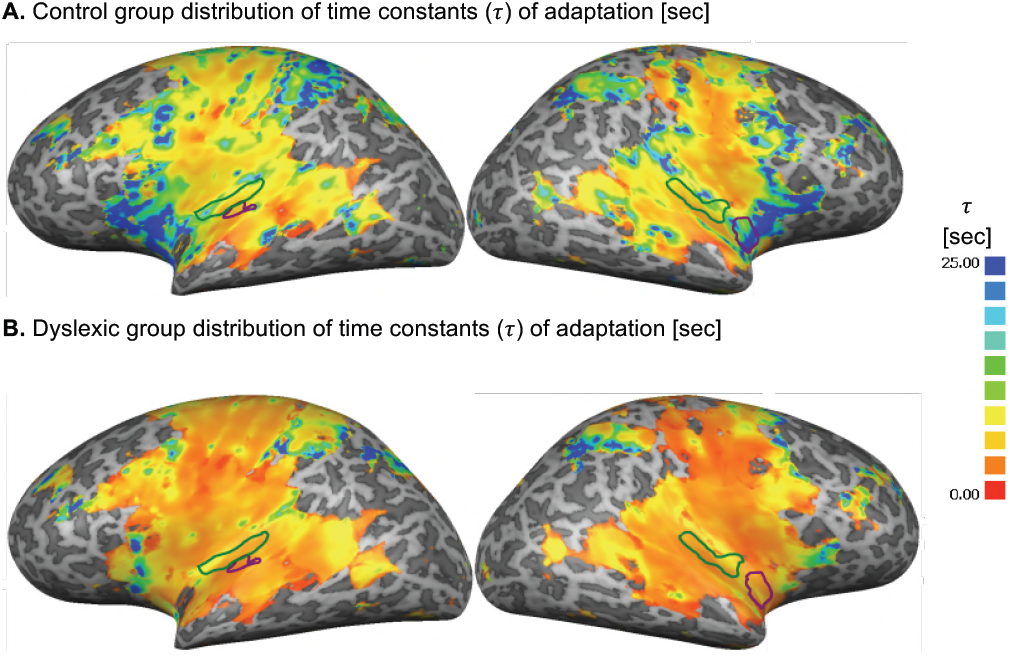
Cortical distribution of the estimated time constants (τ) of adaptation, calculated separately for each of the responding voxels, based on the mean BOLD response. **A.** Controls. **B.** Dyslexics. Dyslexics’ estimated τs were consistently shorter. Significant group differences (Monte-Carlo cluster-level corrected: cluster threshold of 44 voxels) are outlined in purple. Green outlines denote primary auditory cortex ROI.

The whole brain analysis allocated high level areas in the left superior temporal lobe and the right insular cortex. However, it did not allocate a consistent cluster of significant group-difference voxels in the primary auditory cortex (Zatorre, Belin, & Penhune, 2002). To test whether the primary auditory cortex would show a similar group difference when its BOLD response was averaged across voxels, we delineated a ROI in each hemisphere, based on a combined cytoarchitectonic (Morosan et al., 2001) and myeloarchitectonic (Dick et al., 2012) definition (we included the three sub-regions of the primary auditory cortex: Te1.1, Te1.0 and Te1.2). We fitted the exponential decay model to the βs averaged over the right and the left auditory cortices (composed of 99 voxels each, denoted by the green outlines in Figures 1A-B). We found significant differences between the groups’ τs in the left auditory cortex (*z* = 2.6, *p* < 0.01, effect size *r* = 0.42). In the right primary auditory cortex, the difference between controls’ and dyslexics’ τ showed the same trend, but did not reach significance (*z* = 1.5, *p* = 0.15, effect size *r* = 0.23. Mann-Whitney U-tests). Figure 2 shows the βs estimated for the left and right primary auditory cortices of the controls (blue) and dyslexics (red) on each of the four TOA blocks.

**Figure 2.**
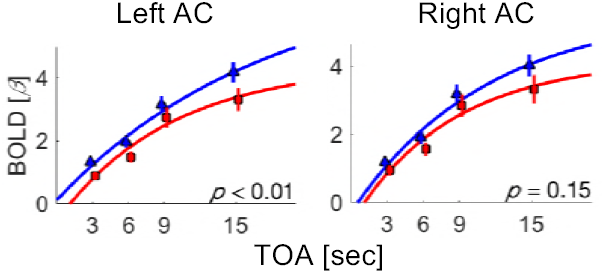
BOLD response as a function of TOA in the primary auditory cortex of each hemisphere. Blue – control. Red – dyslexic. AC – auditory cortex.

Taken together, the whole brain and ROI analyses revealed a significant group difference in the time scales of adaptation in the left superior temporal cortex, the primary auditory cortex, and the right insular cortex. Nevertheless, the general trend of dyslexics’ shorter adaptation was consistent across all responding voxels.

## Discussion

We characterized the cortical distribution of dyslexics’ and controls’ decay of BOLD adaptation, thus extending our previous behavioral and ERP study (Jaffe-Dax et al., 2017). We found a broadly distributed tendency for shorter adaptation in dyslexia. We further assessed group difference in the left and right primary auditory cortices, for which previous reports are mixed. For example, Clark et al. reported early anatomical abnormalities (Clark et al., 2014), whereas Boets et al. (2013) reported adequate stimulus resolution. We found a significant group difference in the left primary auditory cortex, and a similar tendency, which did not reach significance, in the right primary auditory cortex.

The broad distribution of abnormally short adaptation in dyslexia is in line with recent observations of a domain general abnormally small adaptation in dyslexia (Perrachione et al., 2016). Perrachione et al. compared BOLD responses to stimulus repetitions with responses to different auditory and visual stimuli, and found reduced stimulus-specific adaptation in high-level the auditory (superior temporal), visual (fusiform and LO), and associative (insular and inferior frontal) cortices. Their repeated (and non-repeated) stimuli were presented over a similar temporal window as in the current study. Therefore, dyslexics’ abnormally small adaptation may stem from its shorter duration; in other words, dyslexics’ accumulative adaptation across a time window of ˜10 seconds was smaller than controls’ because it largely recovered. In line with the observation of dyslexics’ domain general reduced adaptation, a reduced effect of previous trials was also found behaviorally in the visual modality when performance was measured with serial visual (spatial frequency) discrimination (Jaffe-Dax, Lieder, Biron, & Ahissar, 2016). Together, these studies are consistent with the interpretation that dyslexics’ sensory processing is adequate, but their cortical neural adaptation is abnormally short, yielding shorter implicit memory traces.

## Methods

In the 2-tone frequency discrimination task, subjects were asked to indicate which of two sequentially presented tones had a higher pitch. The tones were 50ms long, presented at comfortable intensity, and were drawn from a uniform distribution between 800-1250 Hz. The frequency difference within each pair was randomly drawn between 1-20% (following the protocol in Jaffe-Dax et al., 2017). In the pre-training session (8 minutes) each participant performed 16 trials of each of the 4 Trial Onset Asynchronies (TOAs) of 3, 6, 9, or 15 seconds, administered in 4 separate blocks in random order. These TOAs are longer than those in our previous ERP experiment (1.5, 3, 6, and 9 seconds, Jaffe-Dax et al., 2017), since the controls’ ERP (N1 and P2) response at 9 seconds was still larger than at 6 seconds. In the scanner, each participant performed 3 runs of 4 blocks of 16 trials (48 trials in each TOA). Each block had a constant TOA of either 3, 6, 9, or 15 seconds. This number of trials was sufficient for estimating τ based on the magnitude of the BOLD response. However, it was too small for robust estimation of behavioral context effects, which are based on the difference in success rate (binary scores for each trial) between trials that gain and those that are hampered by the context (Jaffe-Dax et al., 2017). Stimuli were digitally constructed using Matlab 2015b (The Mathworks Inc., Natwick, MA) and administered through inserted sound attenuating MR compatible S14 earphones (Sensimetrics Corporation, Malden, MA). The demographic, cognitive and reading assessments of this cohort are described in Jaffe-Dax et al., 2017.

Before the functional scan, high-resolution (1 × 1 × 1 mm resolution) T1-weighted magnetization-prepared rapid acquisition gradient-echo (MPRAGE) images were acquired using a 3-T Magnetom Skyra Siemens scanner and a 32-channel head coil at the ELSC Neuroimaging Unit (ENU). The cortical surface was reconstructed from the high-resolution anatomical images using standard procedures implemented by the BrainVoyager QX software package (version 2.84; Brain Innovation, The Netherlands). The functional T2*-weighted MRI protocols were based on a multislice gradient echo-planar imaging and obtained under the following parameters: TR = 1 s, TE = 30 ms, flip angle = 90°, imaging matrix = 64 × 64, field-of-view = 192 mm; 42 slices with 3 mm slice thickness and no gap were oriented in AC-PC plane, covering the whole brain, with functional voxels of 3 × 3 × 3 mm and multiband parallel imaging with an acceleration factor of 3 (Moeller et al., 2010).

Preprocessing of functional scans in BrainVoyager included 3D motion correction, slice scan time correction, and removal of low frequencies up to 3 cycles per scan (linear trend removal and high-pass filtering). The anatomical and functional images were transformed to the Talairach coordinate system using trilinear interpolation. Each voxel’s time course was z-score normalized and smoothed using a 3D Gaussian filter (FWHM of 4 mm). A standard (2 gamma) hemodynamic response function (Friston et al., 1998) was convolved with the trial timings of each TOA block to build four predictors for the subsequent GLM analysis. For all task-responsive voxels (*p* < 0.001, FDR corrected; Benjamini & Yekutieli, 2001), each TOA condition was modeled separately to account for its contribution to the measured BOLD signal in each voxel. Specifically, a single β value was obtained for each TOA condition. An exponential decay model (see Results) was fitted to these β values, and its parameters were estimated for each voxel in each subject using a least-square method. For ROI analysis, the MNI coordinates of auditory cortex subdivision were obtained from Morosan et al. (2001) and translated into Talairach coordinates using Yale BioImage Suite Package (sprout022.sprout.yale.edu/mni2tal/mni2tal.html; Lacadie, Fulbright, Rajeevan, Constable, & Papademetris, 2008). The BOLD signal was averaged for each ROI and then the β values of the four TOA blocks were fitted to the exponential decay.

Non-parametric tests (Mann-Whitney’s U-test) were used for group comparison, since we did not assume a normal distribution (Jaffe-Dax et al., 2017). Whole-brain significance results were corrected for multiple comparison false positive biases by a Monte-Carlo cluster correction (Forman et al., 1995; Goebel, Esposito, & Formisano, 2006).

## Acknowledgments

We thank Udi Zohary, Yuval Porat, Luba Daikhin, Tal Golan and Zvi Roth for their valuable feedback on this manuscript. This study was supported by the Israel Science Foundation (ISF grant no. 616/11 and Canada-Israel grant no. 2425/15), the Gatsby Charitable Foundation, The German-Israeli Foundation for Scientific Research and Development (grant no. I-1303-105.4/2015), Canadian Institutes of Health Research (CIHR), The International Development Research Center (IDRC) and the Azrieli Foundation.

